# *Echis carinatus* snake venom metalloprotease-induced toxicities in mice: therapeutic intervention by a repurposed drug, tetraethylthiuram disulfide (disulfiram)

**DOI:** 10.1101/2020.07.17.208348

**Authors:** Gotravalli V. Rudresha, Amog P. Urs, Vaddarahally N. Manjuprasanna, Mallanayakanakatte D. Milan Gowda, Krishnegowda Jayachandra, Rajesh Rajaiah, Bannikuppe S. Vishwanath

**Author notes:** Corresponding author(s) Prof. B S Vishwanath, Dr. Rajesh Rajaiah.

## Abstract

*Echis carinatus* (EC) is known as saw-scaled viper and it is endemic to the Indian subcontinent. Envenoming by EC represents a major cause of snakebite mortality and morbidity in the Indian subcontinent. Zinc (Zn^++^)-dependent snake venom metalloproteases (SVMPs) present in *Echis carinatus* venom is well known to cause systemic hemorrhage and coagulopathy in experimental animals. An earlier report has shown that ECV activates neutrophils and releases neutrophil extracellular traps (NETs) that blocks blood vessels leading to severe tissue necrosis. However, the direct involvement of SVMPs in the release of NETs is not clear. Here, we investigated the direct involvement of EC SVMPs in observed pathological symptoms in a preclinical setup using zinc (Zn^++^) metal chelator, Tetraethylthiuram disulfide (TTD)/disulfiram. TTD potently antagonizes the activity of SVMPs-mediated ECM protein degradation in vitro and skin hemorrhage in mice. In addition, TTD protected mice from ECV-induced footpad tissue necrosis by reduced expression of citrullinated H3 (citH3) and myeloperoxidase (MPO) in footpad tissue. TTD also neutralized ECV-induced systemic hemorrhage and conferred protection against lethality in mice. Moreover, TTD inhibited ECV-induced NETosis in human neutrophils and decreased the expression of peptidylarginine deiminase (PAD) 4, citH3, MPO and pERK. Further, we demonstrated that ECV-induced NETosis and tissue necrosis is mediated via PAR-1-ERK axis. Overall, our results provide an insight into SVMPs-induced toxicities and the promising protective efficacy of TTD can be extrapolated to treat severe tissue necrosis complementing ASV.

## Introduction

According to the World Health Organization, snakebite is a global public health problem and a neglected tropical disease [1, 2]. Snakebites are often associated with severe local manifestations including inflammation, hemorrhage, blistering, skin damage, coagulopathy and progressive tissue necrosis at the bitten site [3, 4]. These local manifestations are never the less a pathological condition, caused by a mixture of toxins rather than a single toxin present in the venom [4]. Unfortunately, treatment of snake envenomation-induced progressive tissue necrosis persists even after anti-snake venom (ASV) administration [5, 6]. Hence, treating progressive tissue necrosis is still a challenging issue for the existing strategies of snakebite management. In addition, studies on progressive tissue necrosis induced by viper bite have clearly revealed the direct involvement of metzincin family matrix-degrading snake venom metalloproteases (SVMPs)[4, 7, 8] and hyaluronidases (SVHYs) **[9, 10]**.

*E. carinatus* (EC) also known as a saw-scaled viper is the only Echis species found in India **[11]**. Echis bite is responsible for the highest mortality and causes severe local manifestations, due to rich in zinc (Zn^++^)-dependent SVMPs that act on the basement membrane and destabilize the tissue [12, 13]. In addition, ECV activates neutrophils and releases their nuclear and granular contents to the extracellular space leading to neutrophil extracellular traps (NETs) the process is described as NETosis [14]. They showed that NETs released from neutrophils lock the venom toxins and results in blockage of blood vessels leading to tissue necrosis [14]. However, the venom toxin/s responsible for NETs formation and cellular mechanism involved is unclear.

Several studies have demonstrated that SVMPs of ECV are known to contribute to the pathophysiology by causing many local manifestations **[4, 15-17]**. SVMPs have resemblance in catalytic site architecture and structural domains with metzincin family proteases such as MMPs and ADAMs [18]. A few reports showed the activation of MAPKs by MMPs via protease-activated receptor (PAR)-1 [19]. Since SVMPs are catalytically related to MMPs, we hypothesized that EC SVMPs induce NETosis and intracellular signaling cascade via PAR-1. Here, we have demonstrated that EC SVMPs-induced NETosis is mediated via PAR-1-ERK signaling axis, responsible for severe tissue necrosis. In this line, we have used an Antabuse drug, disulfiram/TTD repurposed as therapeutic for ECV-induced toxicities in preclinical setup.

## Materials and methods

### Venom

Lyophilized powder of *Echis carinatus* venom (ECV) was purchased from Irula Snake-Catchers Co-operative Society Ltd., (Chennai, India). The required amount of venom was re-dissolved in PBS and centrifuged at 9000 g for 10 min to remove debris. Aliquots of venom were kept at −20°C until further use. The protein content of venom was determined according to the method of Lowry et al. [20].

### Chemicals and reagents

Tetraethylthiuram (TTD), aristolochic acid (AA), silymarin (SLN), phorbol 12-myristate 13-acetate (PMA), porcine skin gelatin, collagen type 1, laminin, fibronectin, Hoechst stain, DNase 1, ficoll-paque, dextran and phosphatase inhibitor cocktail were obtained from Sigma-Aldrich (Bangalore, India). BSA, ethanol, dimethyl sulfoxide (DMSO; HPLC grade), Tween-20 and Hank’s balanced salt solution (HBSS) were purchased from HiMediaLaboratories, Pvt. Ltd. (Mumbai, India). SCH79797 (PAR-1 antagonist) and GB-83 (PAR-2 antagonist) were purchased from Cayman Chemicals (Michigan, USA). U0126 (MEK 1/2 inhibitor), antibodies against p-ERK and β-actin were purchased from Cell Signaling Technology (Massachusetts, USA). HRP tagged anti-rabbit IgG and anti-mouse IgG were procured from Jackson ImmunoResearch (Philadelphia, USA). The rabbit polyclonal anti-citH3, rabbit polyclonal anti-H3, mouse monoclonal anti-myeloperoxidase (anti-MPO) and anti-PAD4 were obtained from Abcam (Cambridge, UK). All other chemicals and reagents used in this study are analytical grade.

### Animals

Adult Swiss albino mice (6 to 8 week old male or female) weighing 20 - 25 g were obtained from the Central Animal House Facility, Department of Studies in Zoology, University of Mysore, Mysuru, India. The animal experiments were approved by the Institutional Animal Ethical Committee, University of Mysore, Mysuru, India (Approval number: UOM/IAEC/20/2016). During all experiments, animal care and handling were in accordance with the guidelines of the Committee for the Purpose of Control and Supervision of Experiments on Animals (CPCSEA).

### Human plasma

Human blood was drawn from the antecubital veins of healthy adult volunteers who were provided with written informed consent. All the experiments were approved by the Institutional Human Ethical Committee, University of Mysore, Mysuru, India (Approval number: IHEC-UOM No. 120 Ph.D/2015-16), and conducted in accordance with the ethical guidelines.

### Gelatinolytic activity

The gelatinolytic activity was performed by substrate gel assay as described by Heussen and Dowdle, with some modifications [21]. ECV, 5 µg was loaded onto a 10% SDS polyacrylamide gel (SDS-PAGE) impregnated with 0.08% of gelatin. After electrophoresis, SDS was removed by incubating gel with 2.5% Triton X-100 for 1 h, followed by an extensive wash with distilled water. The gel was incubated overnight at 37°C in incubation buffer, 50 mM Tris-HCl, pH 7.6 containing 0.9% NaCl, 10 mM CaCl_2_, 10 mM ZnCl_2_. The gel was stained with Coomassie brilliant blue-G250 (CBB-G250) and a clear zone indicates the gelatinolytic activity of ECV. For inhibition studies ECV was pre-incubated with different concentrations of TTD (1, 5, 10 and 20 mM), AA (10 and 20 mM) and SLN (10 and 20 mM) for 5 min at 37°C and assay was performed as described above.

### ECM protein hydrolyzing activity

ECM protein hydrolyzing activity was performed according to the method of Baramova et al. with slight modifications [22]. ECM proteins, collagen type-I and −IV, laminin and fibronectin (50 µg each) were incubated with 5 µg of ECV, separately in a total reaction volume of 40 µl with Tris-HCl buffer (10 mM; pH-7.6) at 37°C for 3 h. The reaction was terminated by adding 20 µl of reducing sample buffer (4% SDS, 6% β-mercaptoethanol and 1 M urea) and boiled for 5 min. An aliquot of 40 µl of this sample was loaded onto 7.5% SDS-PAGE and electrophoresed. The cleavage pattern of ECM proteins was visualized by staining with CBB-G250. For inhibition studies, similar experiments were carried out by pre-incubating ECV with different concentrations of TTD (1, 5, 10 and 20 mM), AA (10 and 20 mM) and SLN (10 and 20 mM) for 5 min at 37°C and electrophoresed as described above.

### ECV-induced skin hemorrhage in mice

Hemorrhagic activity was performed as described by Kondo et al. with suitable modifications [23]. Mice were injected (i.d.) with 5 µg of ECV and control mice received saline. After 21/2 h, mice were sacrificed and the inner dorsal surface of the skin was surgically removed and photographed, and the hemorrhagic spot was quantitated using a graph sheet. For neutralization studies, various concentrations of TTD (13.25, 26.5 and 53 μg), AA (20 and 40 μg) and SLN (15 and 30 μg) were injected 15 min post ECV injection. Neutralization of hemorrhagic activity was measured in terms of decreased area of the hemorrhagic spot in comparison to ECV-induced hemorrhagic area.

### ECV-induced mice footpad tissue necrosis

ECV-induced mice footpad tissue necrosis was performed as described by Rudresha et al. with suitable modifications [24]. ECV (LT_50_; 2.21 mg/kg body weight) was administered to the mice footpad (n-5; Intraplantar injection). The time for onset of footpad injuries was recorded for each mouse. The severity of the injury was visually judged and scored based on a 5-point scale; 0-no injury, 1-edema with mild hemorrhage, 2-edema with severe hemorrhage causing discoloration of the footpad, 3-edema with severe hemorrhage and necrosis, 4-severe hemorrhage and necrotized footpad, 5-necrotized little toe detached from limb. The footpad injury observations were recorded every day for 8 days after venom injection. To evaluate the protection of ECV-induced tissue necrosis, mice were anesthetized with pentobarbitone (30 mg/kg; i.p.) and TTD (2.15 mg/kg) was administered to the venom injection site 30 min post venom injection. DNase 1 (100 U) was used as a positive control. To assess the effect of SCH79797 (PAR-1 antagonist) on ECV-induced footpad tissue necrosis, SCH79797 (1.5 mg/kg) was administered to the mice footpad 15 min before to the ECV (1/2LT_50_; 1.10 mg/kg) injection. The onset of footpad injuries was recorded as described above.

### ECV-induced mortality and systemic hemorrhage in mice

Lethal toxicity of ECV was determined according to the method of Meier and Theakston, with some modifications [25]. To determine the anti-venom potential of TTD in preventive regimen, ECV (11/2 LT_50_; 3.31 mg/kg) was pre-incubated with TTD (2.15 mg/kg) or effective dose ASV (ED ASV), separately for 5 min at 37°C and injected intra peritoneally (i.p.) to mice. For the treatment model, TTD (2.15 mg/kg) or ED ASV was injected 30 min post of venom administration (i.p.). The signs and symptoms of toxicities were observed up to 24 h and survival time was recorded. For systemic hemorrhage and bleeding studies, mice received ECV (LT_50_; 2.21 mg/kg, i.p) and challenged with TTD (2.15 mg/kg; i.p) or ED ASV, 30 min post ECV injection. After 3 h peritoneal cavity was observed for symptoms and photographed.

### Isolation of neutrophils from human blood

Human neutrophils were isolated from healthy volunteers blood, according to the method of Halverson et al. with slight modifications [26]. The blood was collected and mixed with acid citrate dextrose in a 5:1 volumetric ratio, followed by dextran sedimentation and hypotonic lysis to remove red blood cells. The cell pellet was suspended in 2 ml of PBS and subjected to density gradient centrifugation at 210 g using ficoll-paque for 30 min at 4°C. The neutrophils settled at the bottom as a cell pellet were washed twice with PBS and centrifuged at 210 g and re-suspended in HBSS. The cells were counted using the Neubauer hemocytometer and the required cell density was adjusted using HBSS.

### Analysis of NETs formationand its markers in human neutrophils

NETs formation was analyzed using Hoechst stain according to the method of Katkar et al. with suitable modifications [14]. Human neutrophils (2×10^5^/ml) were seeded on 13 mm round coverslips placed in 12 well plates in 500 µl of HBSS-1 and allowed cells to adhere to the coverslips for 30 min at 37°C in the presence of 5% CO_2_. Then, the cells were stimulated with different concentrations of ECV (5, 10, 25 and 50 µg/ml) for 21/2 h. After incubation, cells were fixed with 4% paraformaldehyde for 30 min and stained with Hoechst stain (1:10,000) for 15 min. Cells were analyzed for NETs formation and images were acquired on a BA410 fluorescence microscope (Motic) attached to a DS-Qi2 monochrome CMOS sensor camera. The NETs percentage was calculated in 5 non-overlapping fields per coverslip. For inhibition studies, similar experiments were carried out by pre-incubating ECV (25 µg) with different concentrations of TTD (1, 5, 10 and 20 mM), AA (10 and 20 mM) and SLN (10 and 20 mM) for 5 min at 37°C and NETs percentage was calculated as described above.

For the analysis of ECV-induced citH3, PAD4 and MPO expression, neutrophils were treated with ECV (25 µg) that was pre-treated with various concentrations of TTD (1, 5, 10 and 20 mM), AA (10 and 20 mM) and SLN (10 and 20 mM) at 37°C for 5 min. After 21/2 h, neutrophils were washed and lysed with lysis buffer containing PMSF and phosphatase inhibitor cocktail and stored at − 20°C overnight. Lysates were centrifuged at 9000 g for 10 min and tested for the expression of protein using Western blotting.

To test the effect of pharmacological inhibitors on ECV-induced NETs and its markers, neutrophils were pre-sensitized without or with U0126 (MEK 1/2 inhibitor), SCH79797 (PAR-1antagonist) and GB-83(PAR-2 antagonist) for 15 min. Then, neutrophils were stimulated with 25 μg of ECV for 21/2 h at 37°C, and NETs were quantitated using Hoechst stain. Cell lysates were used to test the expression of citH3, PAD4, MPO and ERK activation using Western blotting.

### Western blotting

ECV injected mice footpads were excised from respective days 1 to 4 and stored at −20°C. The excised tissue samples were sonicated in cold lysis buffer containing EDTA free protease inhibitor mixture for 30 seconds on ice bath (five passes of 10 seconds) with a 3.0 mm probe sonicator. Tissue homogenates were centrifuged at 9000 g for 10 min and the supernatant was used for protein quantification. Western blotting was performed as described by Rajaiah et al. with suitable modifications [27]. Briefly, an equal amount of proteins (25 - 40 µg) were resolved on SDS-PAGE in Laemmli buffer and electro-blotted onto polyvinylidene difluoride (PVDF) membranes. After blocking for 2 h in TBST containing 5% fat-free milk, membranes were probed against anti-citH3 or anti-PAD4 or anti-MPO or anti-p-ERK antibodies overnight at 4°C according to the manufacturer’s instructions (Abcam, UK and Massachusetts, USA). Membranes were washed extensively (4x) using TBST and incubated with HRP-conjugated anti-rabbit/mouse IgG (1:10,000) for 1 h at room temperature. The blots were developed with an enhanced chemiluminescence substrate for visualization (Alliance 2.7, Uvitec) and bands were quantitated using Image J software. H3 and β-actin were used as the loading control.

### Statistics

The data are represented as the mean ± SEM. The comparison between the two groups was analyzed using the Mann-Whitney U test. The analysis of variance (ANOVA) followed by Bonferroni post hoc test was used for more than two groups using Graph Pad Prism version 5.03 (La Jolla, USA). The comparison between the groups was considered significant if p< 0.05.

## Results

### TTD inhibits ECV-induced ECM protein degradation and hemorrhage

ECV is known to induce the degradation of ECMproteins such as collagen, gelatin, laminin and fibronectin, resulting in hemorrhage at the site of injection [4, 8, 17]. Multiple scientific reports have demonstrated the direct involvement of SVMPs in disrupting the tissue architecture by degrading ECM proteins [7, 28, 29]. Hemorrhagic SVMPs act at the basement membrane and disrupt the capillary wall that results in extravasation [7, 28, 30]. Further experimental evidence suggests that the onset of micro-vessel damage is mediated by the degradation of type IV collagen by the action of SVMPs [7, 30]. Previously, we have shown the neutralizing abilities of zinc-specific chelators against the snake venom-induced progressive tissue damage [31]. Very recently, Albulescu et al. demonstrated the therapeutic intervention of repurposed drug, 2,3-dimercapto-1-propanesulfonic acid for hemotoxic snakebite [32]. In this view, we tested the inhibitory efficacy of an Antabuse drug with chelating property, TTD on ECV-induced ECM protein degradation and hemorrhage in mice, and compared with pharmacological inhibitors of PLA_2_ (AA) and SVHYs (SLN). To begin with, the effect of TTD on ECV-induced gelatinolytic activity was demonstrated by gelatin substrate gel assay. ECV-induced gelatinolytic activity was inhibited by TTD in a concentration-dependent manner **(Fig. 1A)** and maximum inhibition was observed at 20 mM of TTD **(Fig. 1B)**. In addition, TTD also inhibited ECV-induced ECM proteins, collagen, laminin and fibronectin degradation in a concentration-dependent manner **(Fig. 1C - 1F)**. Furthermore, TTD was tested for its action on ECV-induced hemorrhagein mice skin. TTD efficiently neutralized ECV-induced hemorrhage at a tested dose even after 15 min post ECV injection **(Fig. 1G and 1H)**. On the other hand, the snake venom PLA_2_ and SVHYs inhibitors, aristolochic acid (AA) and silymarin (SLN) failed to inhibit ECV-induced ECM protein degradation **(Fig. S1A - S1F)** and hemorrhagic activity **(Fig. S1G and S1H)**.

**Fig 1:**
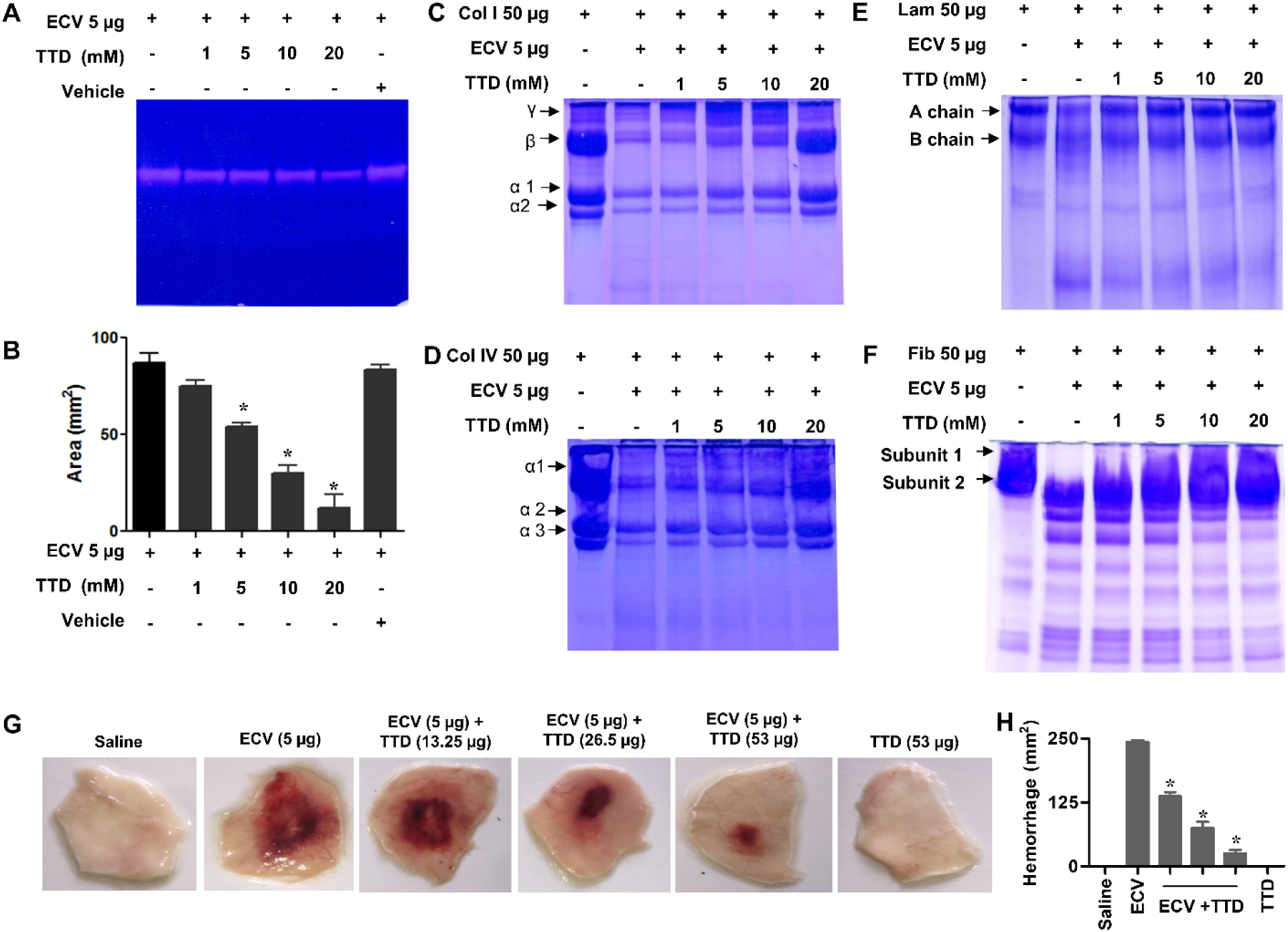
Effect of TTD on ECV-induced ECM protein degradation in vitro and hemorrhage in mice. ECV was pre-incubated without or with various concentrations of TTD at 37°C for 5 min and subjected for gelatinolytic, collagenolytic, laminin and fibronectin hydrolyzing activity. The gelatinolytic activity was performed by gelatin zymogram using 10% gel impregnated with 0.08% gelatin. Clear zones in the gel indicate the hydrolysis of gelatin by ECV **(A)**. The area of gelatinolytic activity was quantitated using a graph sheet represented as area (mm^2^) **(B)**. The data represented as mean ± SEM. *p < 0.05, when compared ECV versus ECV + TTD. For collagen, laminin and fibronectin degradation analysis, the pre-incubated reaction mixture of ECV and TTD was incubated with 50 µg of collagen (Col) type I **(C)**, type IV **(D)**, laminin (Lam) **(E)** and fibronectin (Fib) **(F)** for 3 h at 37°C. The hydrolyzing pattern was analyzed using 7.5% SDS-PAGE and visualized by staining with CBB-G250. Data are representative of two independent experiments. For skin hemorrhage, mice were injected (i.d) with 5 µg of ECV followed by the injection of different concentrations of TTD post 30 min at the site of ECV injection. After 180 min, dorsal patches of mice skin were photographed **(G)** and the hemorrhagic area was measured using graph sheets represented as area (mm^2^) **(H)**. The data represented as mean ± SEM. *p < 0.05, when compared ECV versus ECV + TTD.

### TTD protects ECV-induced mice footpad tissue necrosis with decreased expression of citrullinated H3 (citH3) and myeloperoxidase (MPO)

With promising results of in vitro inhibition of ECV-induced ECM proteins degradation and skin hemorrhage in mice skin, TTD was tested for the neutralization of ECV-induced tissue necrosis using mice footpad model. ECV injection to mice footpad resulted in progressive tissue necrosis and, TTD administration neutralizes ECV-induced tissue injury and restores the normal footpad morphology **(Fig. 2A and 2B)**. Recently, Katkar et al. reported that infiltrated neutrophils to the site of venom injection release chromatin content to the extracellular space as NETs that is responsible for local tissue necrosis [14]. Furthermore, Katkar et al. and Rudresha et al. demonstrated that the intervention of DNase 1 and plant DNase at a right time protected ECV-induced tissue necrosis [14, 33]. In addition, the excessive production of MPO, and citH3 by the action of PAD4 has shown to be crucial for ECV-induced local tissue damage **[14]**. Similar to the previous study, ECV induced the expression of MPO and citH3, and it was efficiently inhibited by TTD **(Fig. 2C - 2E)**. The inhibitory action of TTD on ECV-induced mice footpad necrosis and the expression of MPO and citH3 are more efficient and comparable with DNase 1 **(Fig. 2)**. These data suggested that TTD, which is a well-known chelating agent that neutralizes ECV-induced toxicities via inhibition of SVMPs.

**Fig 2:**
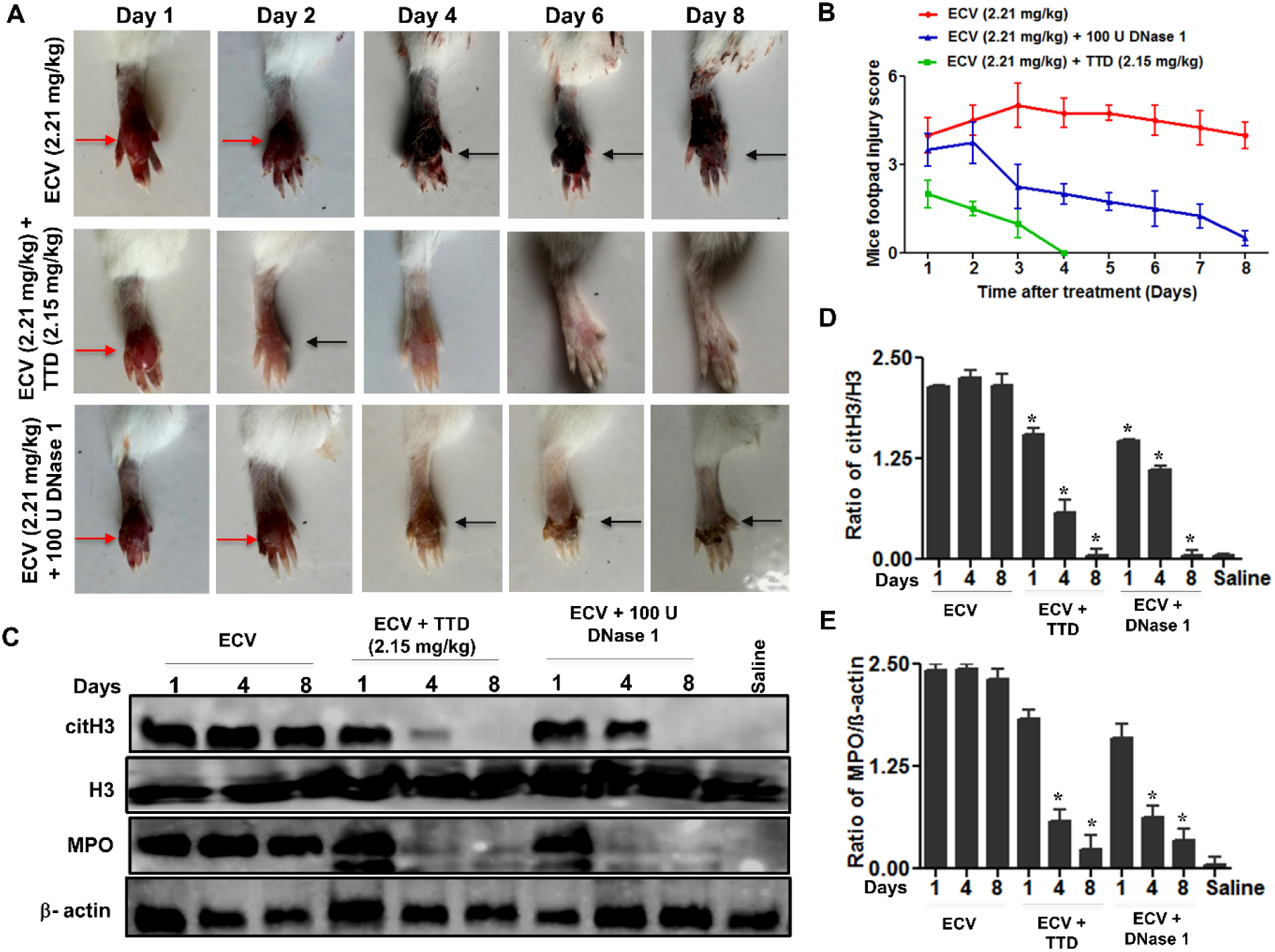
Neutralization of ECV-induced mice footpad tissue necrosis by TTD. Mice footpads were injected with ECV (2.21 mg/kg). After 30 min, mice received either TTD or DNase 1 at the site of venom injection and footpads were photographed from day 1 to day 8 **(A)**. Red arrow indicates edema and black arrow indicates tissue necrosis. ECV-induced footpad injury was measured manually on a scale of 1 to 5 **(B)**. The level of ECV-induced citH3 and MPO in mouse footpad tissue in the absence or presence of either TTD or DNase 1 was analyzed by Western blotting **(C)** and quantitated using H3 and β-actin as a loading control for citH3 **(D)** and MPO **(E)**, respectively. The data represented as mean ± SEM. *p < 0.05, when compared ECV versus ECV + TTD and ECV versus ECV + DNase 1.

### TTD protects mice from ECV-induced lethality and neutralizes systemic hemorrhage

In addition to the induction of progressive tissue necrosis, ECVis lethal when injected at 3.31 mg/kg body weight, and the average survival time is approximately 8 ± 2 h. Since TTD efficiently neutralized ECV-induced tissue necrosis and hemorrhage, its effect on ECV-induced mortality in mice was tested. TTD neutralized ECV-induced lethality and protected mice in both pre-incubation and therapeutic mode (30 min post venom injection) **(Fig. 3A and 3B)**. The protective effect of TTD was comparable to ED ASV both in pre-incubation and therapeutic regimens **(Fig. 3A and 3B)**. ECV is well-known for hemotoxic effect and its envenomation makes blood in-coagulable that leads to the systemic bleeding with disseminated intravascular coagulation [34]. In fact, ECV injection to mouse peritoneum caused severe bleeding and extravasation throughout the peritoneum **(Fig. 3C)**. As TTD protected mice from ECV-induced lethality, it neutralized ECV-induced bleeding in peritoneum even after 30 min post ECV injection and it was comparable with ED ASV as shown in **Fig. 3C**. This indicates that TTD is a potential drug candidate that complements ASV during EC bite.

**Fig 3:**
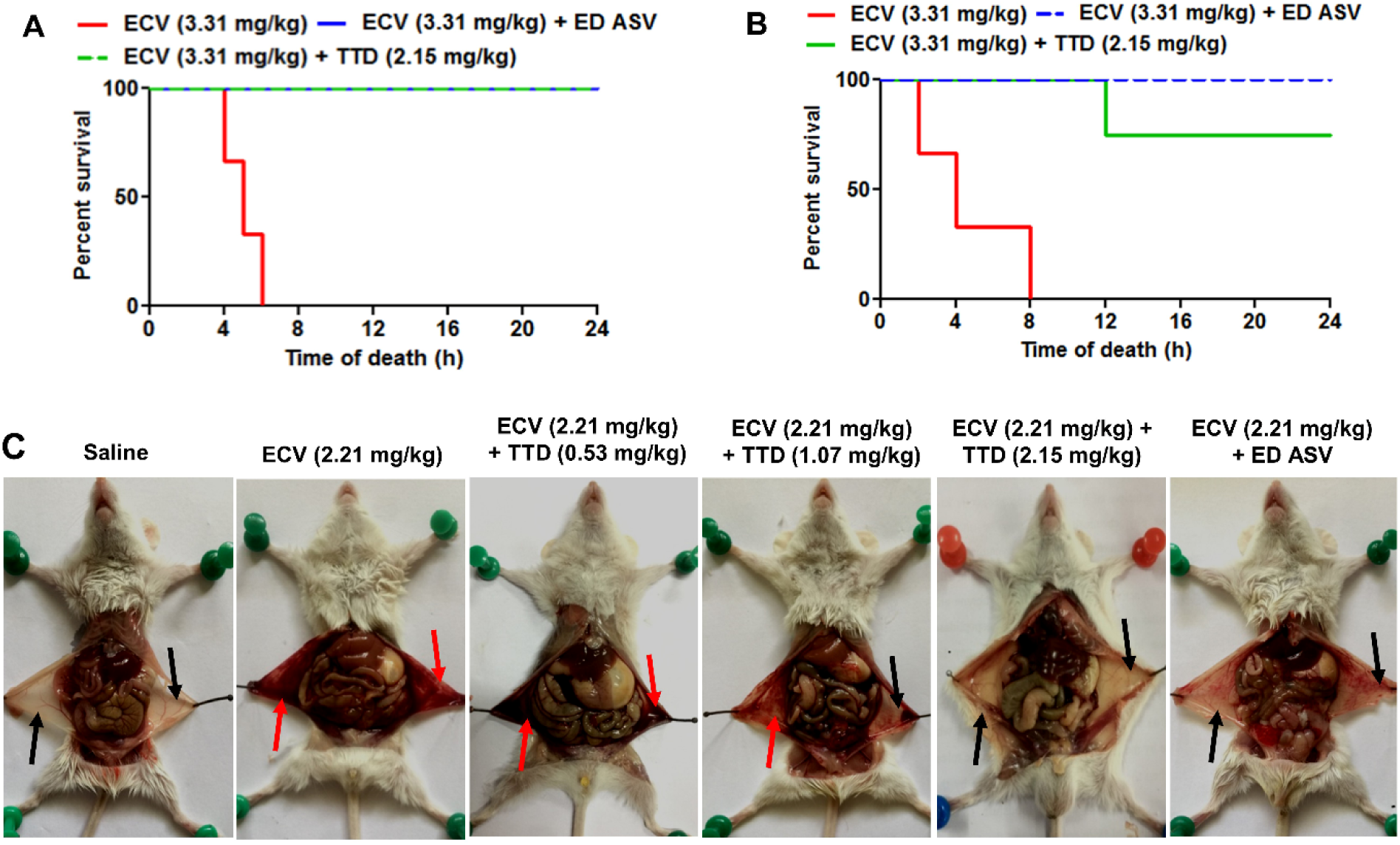
Protection of mice against ECV-induced mortality and systemic hemorrhage by TTD. A lethal dose of ECV (3.31 mg/kg) was pre-treated with TTD (2.15 mg/kg) or an effective dose of anti-snake venom (ED ASV) for 5 min at 37°C and injected (i.p.) to mice. The time taken for mice mortality was recorded for 24 h and graph plotted as percent survival against the time of death **(A)**. In the treatment model, mice received either TTD (2.15 mg/kg) or ED ASV, 30 min post ECV injection (i.p.) and the survival time was recorded for 24 h **(B)**. For the neutralization of systemic hemorrhage, mice received (i.p.) various concentrations of either TTD or ED ASV, 30 min post ECV (2.21 mg/kg) injection (i.p.). Mice were sacrificed after 2 h and peritonea were photographed **(C)**. Red arrow indicates the hemorrhage in the peritoneum cavity and black arrow indicated reduced hemorrhage in the peritoneum. Data are representative of two independent experiments.

### TTD inhibits ECV-induced NETs formation and activation of intracellular signaling in human neutrophils

Neutrophils are the first line innate immune cells recruited to sites of acute inflammation in response to chemotactic signals produced by injured tissue and tissue-resident macrophages [35, 36]. During infection, neutrophils undergo degranulation and ultimately release chromatin as NETs that contribute to the killing of extracellular pathogens [37]. Previously, Setubal et al. demonstrated *Bothrops bilineata* venom in the activation of neutrophils and the release of NETs [38]. Recently, Katkar et al. reportedthe discharged chromatin (NETs) upon ECV treatment is responsible for ECV-induced local tissue necrosis [14]. Similar to the previous reports, we observed ECV-induced chromatin discharge from human neutrophils in a concentration-dependent manner and it was effectively inhibited by TTD **(Fig. 4A and S2A)**. On the other hand, the SVPLA_2_ and SVHYs inhibitors, AA and SLN are failed to inhibit the ECV-induced NETosis **(Fig. S3A and S3B)**. Moreover, ECV treated neutrophils showed increased expression of PAD4, citH3, and MPO and activation of ERK **(Fig. 4B)**. The importance of PAD4 in DNA decondensation by citH3 and DNA expulsion in both mouse and human neutrophils is well documented [39]. Furthermore, TTD significantly reduced ECV-induced NETosis and decreased the expression of PAD4, citH3, and MPO as well as activation of ERK in neutrophils **(Fig. 4A and 4B)**. TTD is a chelating agent that is known to inhibit SVMPs; therefore, these data clearly suggest that SVMPs are directly involved in the activation of ERK and NETs formation.

**Fig 4:**
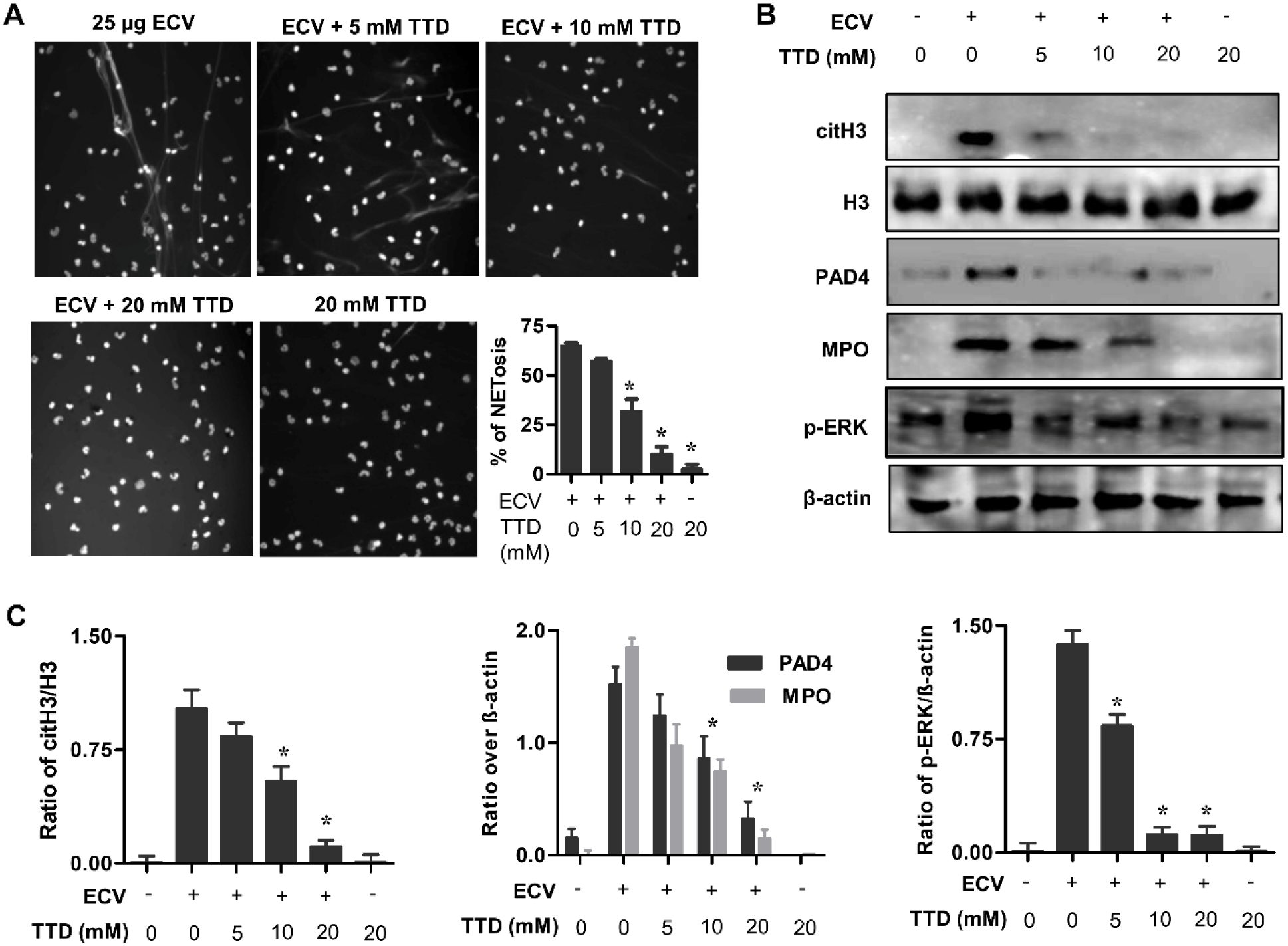
Inhibition of ECV-induced NETs formation by TTD. Human neutrophils were stimulated with ECV (25 µg) pre-incubated (5 min) without or with different concentrations of TTD for 180 min and NETs formation was observed and quantitated **(A)**. ECV-induced citH3, PAD4and MPO in neutrophil cell lysates were analyzed using Western blotting **(B)**. Bands were quantitated using H3 as loading control for citH3 and PAD4 and β-actin as a loading control for MPO **(C)**. The data represented as mean ± SEM. *p < 0.05, when compared ECV versus ECV + TTD.

### ECV-induced NETs formation and tissue necrosis via PAR-1-ERK-mediated axis

PARs are members of the G-protein coupled receptor and cleavage of the extracellular segment, leads to a diverse array of physiological functions [19]. Relatively MMPs are acted on non-canonical sites on the PAR-1 to activate MAPKs pathway to exert their cellular functions [19, 40]. Since MMPs and SVMPs are having structural homology in their catalytic site, we speculated that EC SVMPs activates ERK and NETs formation through PAR-1. To confirm whether ECV induces NETs formation via the PARs, we used PAR-1- and PAR-2 specific antagonists, SCH79797 and GB-83, respectively. ECV activated NETs formation was inhibited in the presence of SCH79797 and not GB-83, suggesting that ECV-induced NETosis is mediated via PAR-1 in human neutrophils **(Fig. 5A and 5B)**. Further, ECV-induced expression of PAD4, CitH3 and activation of ERK was inhibited by SCH79797 **(Fig. 5C)**. On the other hand, MEK inhibitor, U0126 showed a partial effect on ECV-induced NETs formation and the expression of PAD4 and citH3 **(Fig. 5C)**. In support of *in vitro* results in human neutrophils, PAR-1 antagonist neutralized ECV-induced mice footpad tissue necrosis **(Fig. 6A and 6B)**. Overall, these data confirmed the involvement of EC SVMPs-induced tissue necrosis by inducing NETosis and activation of intracellular signalingvia PAR-1 **(Fig. 7)**.

**Fig 5:**
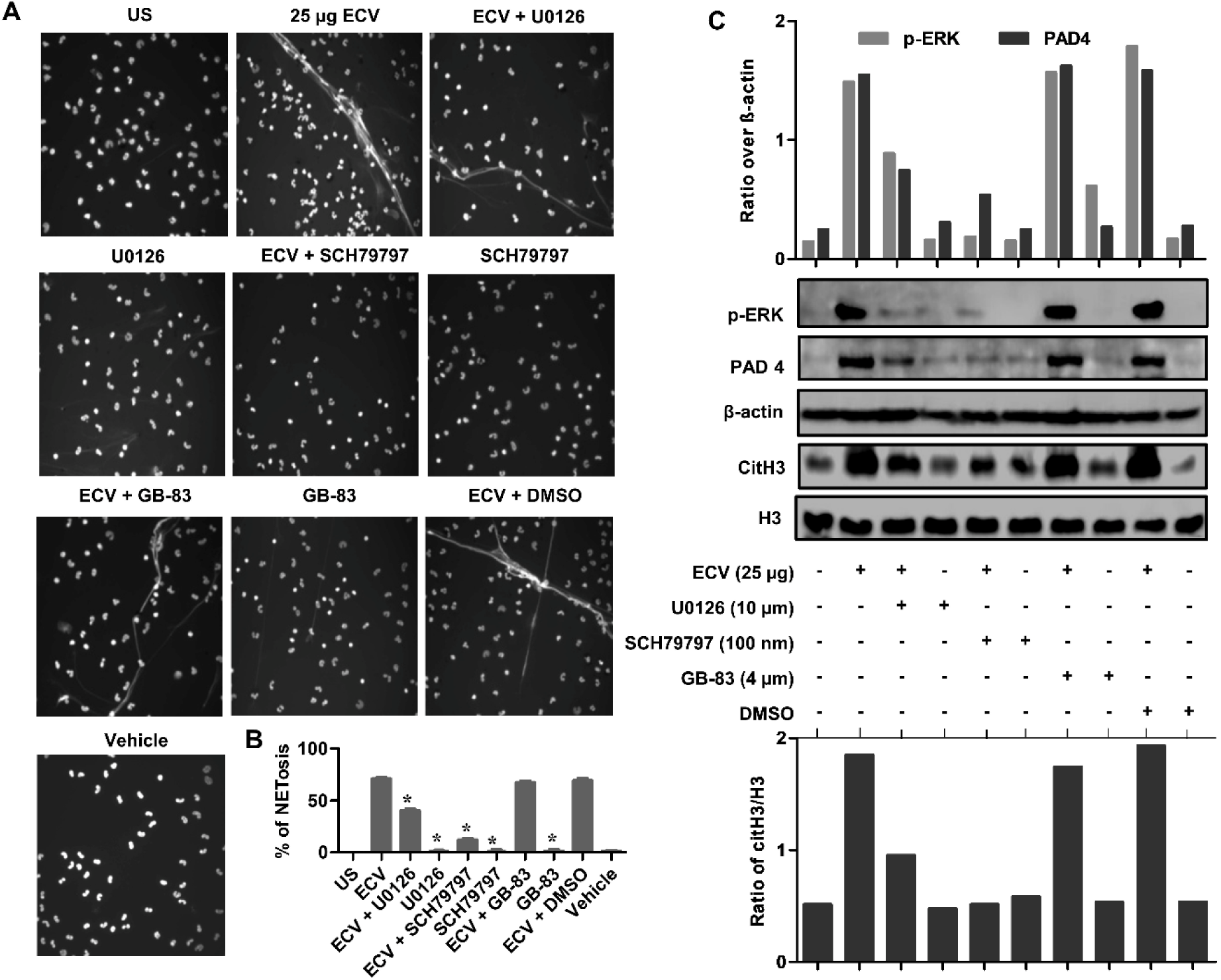
Effect of selective antagonists of ERK and PARs on ECV-induced NETosis and tissue necrosis. Neutrophils were pre-sensitized with selective antagonists of ERK (U0126), PAR-1 (SCH79797) and PAR-2 (GB-83) for 15 min, separately. Pre-sensitized neutrophils were stimulated with 25 µg of ECV for 180 min and cells were stained with Hoechst stain. Neutrophils were photographed under a microscope **(A)** and, NETs were quantitated and represented as percent NETosis **(B)**. The data represented as mean ± SEM. *p < 0.05, when compared ECV versus ECV + antagonists. The whole cell lysates were analyzed for the phosphorylated ERK, expression of PAD4 and citH3 using Western blotting **(C)**. The p-ERK and PAD4 bands were quantitated using β-actin as a loading control. The citH3 bands were quantitated using H3 as a loading control. Data are representative of two independent experiments.

**Fig 6:**
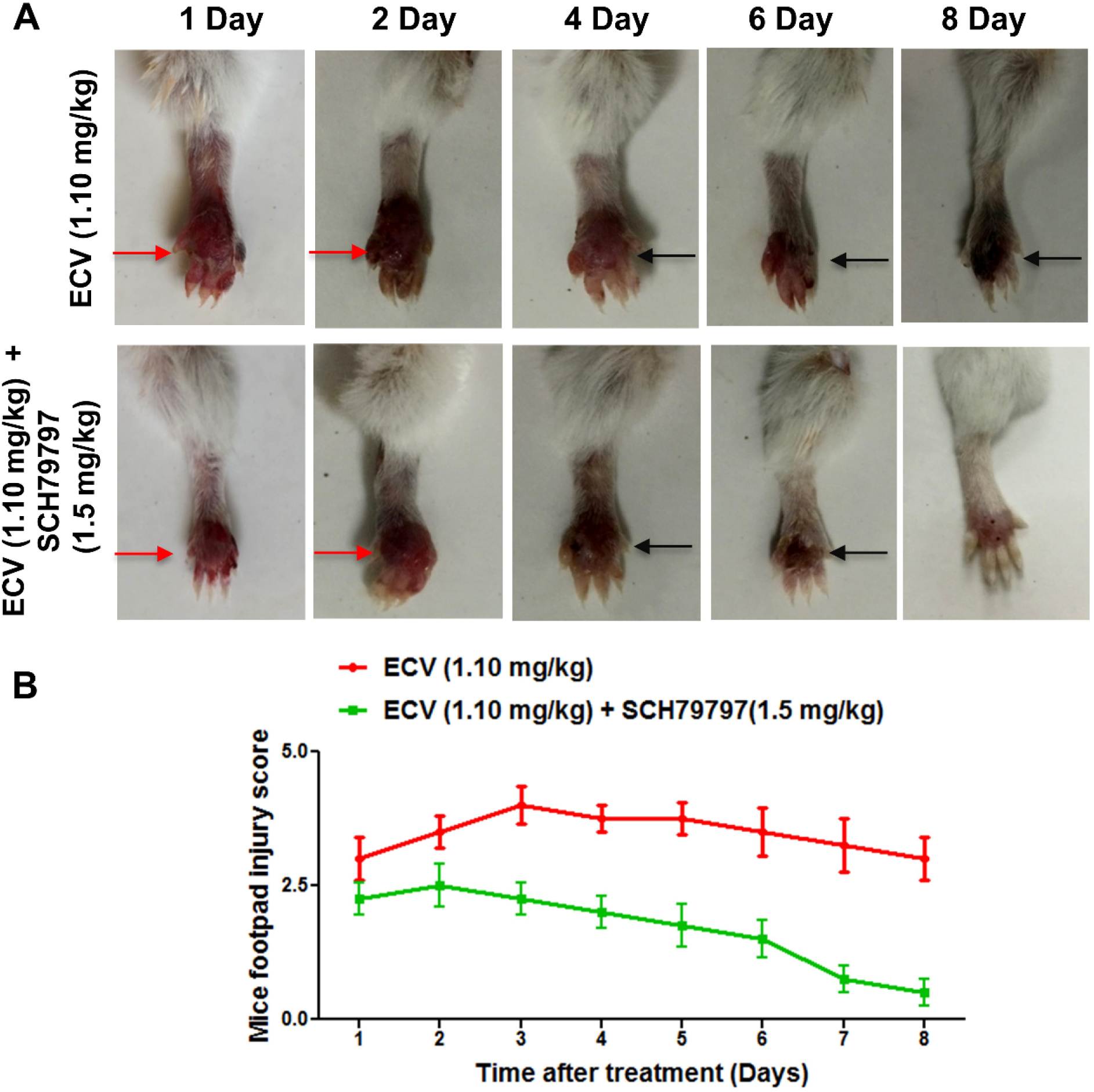
Effect of the selective antagonist of PAR-1 on ECV-induced tissue necrosis. Mice footpads were pre-treated without or with PAR-1 antagonist (SCH79797) for 15 min and followed by the injection of ECV (1.10 mg/kg). Mice footpads were photographed from day 1 to day 8 **(A)** and tissue injury was measured manually on a scale of 1 to 5 **(B)**. Red arrow indicates edema and black arrow indicates tissue necrosis. Data are representative of two independent experiments.

**Fig 7:**
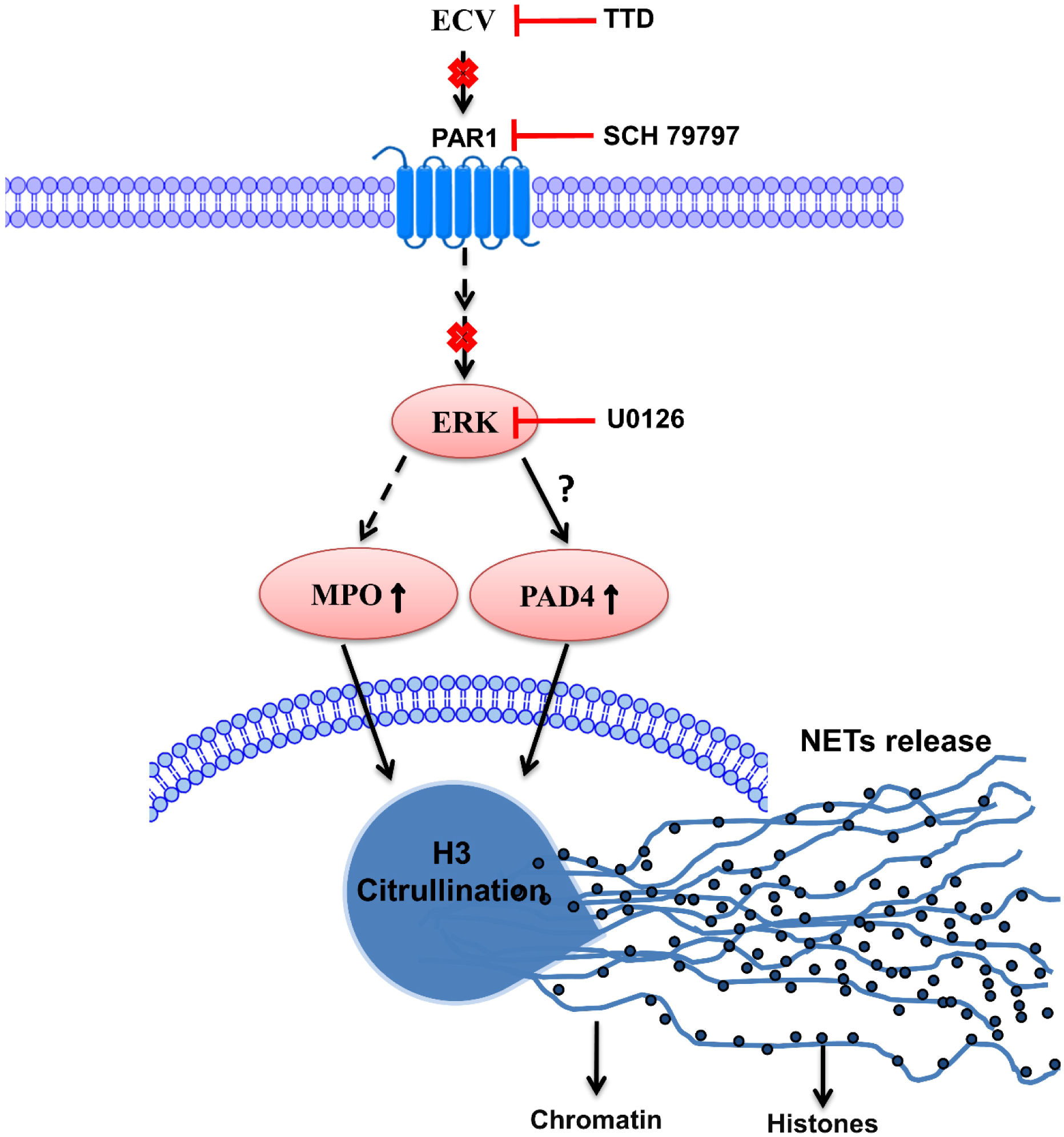
Schematic representation of TTD and pharmacological inhibitors site of action on ECV-induced toxicities. ECV-induced PAR-1-mediated ERK activation might be responsible for increased expression of PAD4, histone citrullination and MPO expressions that are accountable for severe tissue necrosis.TTD and pharmacological inhibitors interfere in ECV-induced signaling/tissue necrosis by inhibiting NETosis and chromatin release.

## Discussion

Viper bites predominantly cause severe local tissue necrosis at the bitten site [41]. Apart from local toxicities, viper bite also exerts their systemic complications such as systemic hemorrhage and coagulopathy leading to hypoxia followed by multi-organ failure and death [32, 42, 43]. SVMPs are one of the major toxins in most of the viper venoms including ECV and they primarily act on ECM components and are responsible for hemorrhagic activity [7, 8, 29-31]. The progressive tissue necrosis induced by viper bites mainly attributed to SVMPs, particularly P-III class metalloproteases [8]. In addition, SVMPs are hemotoxic in nature and interfere with the hemostatic system in snake bite victims [44]. SVMPs are closely related to a disintegrin and metalloproteinases (ADAMs), thus they are also referred to as snake venom ADAMs [45, 46]. Generally, SVMPs contain disintegrin-like (D), cysteine-rich (C), metalloproteinase (M) domain and ADAMDEC-1 that harbors putative zinc (Zn^++^) binding sequence [18, 46, 47]. The secondary structure of the M domain/catalytic domain is topologically similar to matrix metalloproteinases (MMPs) [18, 46]. The catalytic activity of SVMPs is restricted to the M domain and Zn^++^on the ADAMDEC-1 plays a crucial role in their enzymatic activities [18, 46]. Hence, employing specific Zn^++^ chelating agents will be a better approach to counteract the debilitating action of SVMPs. In addition, chelation of Zn^++^ metal ion from SVMPs, by specific Zn^++^chelators rather than anti-venom like molecule is more effective in the management of local tissue necrosis induced by viper venoms.

TTD was the first drug to treat chronic alcoholism and was approved by the FDA 1951 [48]. Since then, many studies have shown repurposing of TTD to treat diverse types of malignant tumors including breast cancer, glioblastoma and pancreatic carcinoma [49]. TTD has also shown therapeutic potential in treating AIDS patients and it is found to be beneficial in treating Lyme disease [50, 51]. Very recently, the intervention of TTD in normalizing body weight in obese mice has been reported [52]. Besides, TTD inhibited MMP-2 and MMP-9 activity by directly interacting with them via a Zn^++^ chelating mechanism [53]. Recently, Albulescu et al. showed that 2,3-dimercapto-1-propanesulfonic acid, a derivative of dimercaprol effectively antagonizes the activity of Zn^++^-dependent SVMPs in vitro and neutralized saw-scaled viper venoms in mice [32]. Previously, we have reported the inhibitory potential of Zn^++^-specific chelating agents; N,N,N’,N’-tetrakis (2-pyridylmethyl) ethane-1,2-diamine, diethylene triaminepenta acetic acid, TTD on ECV-induced toxicities [31]. In sight of these, we demonstrated that Zn^++^ chelating agent, TTD an Antabuse drug can be likely repurposed as a therapeutic candidate in treating ECV-induced toxicities that are mediated by SVMPs.

The proficient hydrolysis of the basement membrane by SVMPs surrounding the blood vessels leads to immediate events of hemorrhage at the site of venom injection [29, 54]. The progression of hemorrhage resulting in localized myonecrosis is due to extensive degradation of structural proteins and severe inflammation [38, 55]. Initially, TTD successfully inhibited ECV-induced degradation of ECM proteins in a dose-dependent manner and also neutralizes the hemorrhage induced by ECV upon challenging studies **(Fig. 1)**. On the other hand, AA and SLN inhibitors failed to inhibit the action of ECV-induced ECM protein degradation and hemorrhage. In support of the neutralization of hemorrhage, TTD treatment could efficiently protect mice footpad from ECV-induced necrosis **(Fig. 2)**. The successful neutralization of ECV-induced ECM proteins degradation and hemorrhage by TTD indicates that SVMPs are the main toxins responsible for ECV-induced toxicities. Further, EC SVMPs are also hemotoxic and interfere in hemostasis by hydrolyzing clotting factors that lead to persistent coagulopathy and death [56]. Most SVMPs are α and β fibrinogenases that act on fibrinogen and making them truncated, and non-functional [56].

A few scientific reports have shown that inhibitors of SVMPs effectively protect mice from viperid snake venom-induced lethality **[31, 32]**. Similarly, TTD was effective in protecting mice from ECV-induced lethality and systemic hemorrhage. These data clearly indicate that TTD has a beneficial effect on neutralizing ECV-induced toxicities in mice.

Neutrophils are the first-line defense immune cells and efficiently arrest pathogens by NETosis at the site of infection [37, 57]. Porto et al. demonstrated the infiltration of neutrophils at the site of viper venom injection [58]. However, the importance of NETosis in ECV-induced toxicities was not clear until Katkar et al. reported the critical role of NETosis in ECV-induced local tissue damage [14]. NETosis results in the blockage of blood vessels preventing venom from entering into the circulation. The accumulated venom-NETs complexes at the site of venom injection lead to the progressive tissue necrosis [14]. In addition, NETosis in non-healing wounds is noticeable by increased expression of PAD4, citH3 and MPO level [14, 59]. However, the previous study did not explain in the context of the toxin that is responsible for ECV-induced NETosis and toxicities [14]. The inhibition of ECV-induced NETosis and reduced levels of PAD4, citH3 and MPO expression by TTD confirms the direct involvement of EC SVMPs in the induction of NETosis.

Nonetheless, the neutralized ECV-induced tissue necrosis and systemic hemorrhage by TTD correlated with the decreased ECV-induced NETosis. However, the mechanism of how ECV/SVMPs induce NETosis and toxicities is largely unknown. There are multiple scientific reports suggesting that the MMPs exert their effects by cleaving PARs and play an important role in vascular functions **[19, 40]**. Moreover, MMPs bind and cleave the extracellular N terminus of PAR-1 to release a tethered ligand and activate the intracellular G proteins across the membrane and initiate intracellular signaling cascade [19, 60]. The inhibition of MMP-1 induced PAR-1 cleavage restricts the activation of MAPKs [61]. SVMPs belong to metzincin super-family and they are known to activate MAPKs signaling pathways in immune cells which results in elevated levels of pro-inflammatory mediators such as TNF-α, IL-1β and IL-6 leading to chronic inflammation [62]. Similarly, EC SVMPs mediates the phosphorylation of ERK in human neutrophils and it was completely inhibited by TTD **(Fig. 4)**. Similar to MMP-1, EC SVMPs might cleave PAR-1 at the non-canonical site and activate downstream MAPKs signaling. Finally, ECV-induced NETosis and tissue necrosis in experimental animals are effectively neutralized by PAR-1 antagonists **(Fig. 5 and 6)**. Overall, current findings indicate that direct involvement of PAR-1 and downstream MAPKs signaling cascade in EC SVMPs-induced toxicities in mice.

## Acknowledgments

Authors thank UGC (MRP-MAJOR-BIOC-2013-12157) and DST-SERB (EEQ/2017/000737) for financial assistance. Rudresha G V [No.F4-1/2006(BSR)/7-366/2012 (BSR) dated 22-10-2013] and Amog P Urs [Ref. No: 22/12/2013(ii) EU-V] thank University Grants Commission(UGC), New Delhi and Manjuprasanna V N [DV2/30/PDF/PA/IOE/2010-11(Vol-II)] thank Institute Of Excellence [Funded by Ministry of Human Resource Development (MHRD), Government of India], University of Mysore, Mysuru for fellowships. Authors thank Central Animal Facility, University of Mysore, for providing animals. Authors thank Prof. Manjunath Kini R, Dr. Nanjaraj Urs A N and Dr. Suvilesh K N for providing valuable suggestions during the project. Authors also thank Mr. Sumanth M S and Mr. Abhilasha K V for their help during experiments and neutrophils isolation.

## Abbreviations

ECV: *Echis carinatus* venom
ECM: Extracellular matrix
MPO: Myeloperoxidase
SVMPs: Snake venom metalloproteases
SVHYs: Snake venom hyaluronidases
MMPs: Matrix metalloproteases
TNF α: Tumor necrosis factor alpha
IL 6: Interleukin 6
NETosis: Neutrophil extracellular traps
TTD: Tetraethylthiuram disulfide/disulfiram
CitH3: Citrullinated H3
PAD4: Peptidylarginine deiminase 4
PAR: Proteases activated receptor

